# Searching for replicable associations between cortical thickness and psychometric variables in healthy adults: empirical facts

**DOI:** 10.1101/2020.01.10.901181

**Authors:** Shahrzad Kharabian Masouleh, Simon B. Eickhoff, Sarah Genon

**Author notes:** Author’s email addresses. Corresponding authors: Shahrzad Kharabian Masouleh, PhD Post-doctoral researcher Institute of Neuroscience and Medicine, Brain & Behaviour (INM-7) Research Centre Jülich, Germany E-Mail; Sarah Genon, PhD Research Group Leader Cognitive Neuroinformatics Institute of Neuroscience and Medicine, Brain & Behaviour (INM-7) Research Centre Jülich, Germany.

## Abstract

The study of associations between inter-individual differences in brain structure and behaviour has a long history in psychology and neuroscience. Many associations between psychometric data, particularly intelligence and personality measures and local variations of brain cortical thickness (CT) have been reported. While the impact of such findings often go beyond scientific communities, resonating in the public mind, their replicability is rarely evidenced.

Here, we empirically investigated the replicability of associations of an extended range of psychometric variables and CT in a large cohort of healthy adults. Our analyses revealed low likelihood of significant associations. Furthermore, significant associations from exploratory analyses showed overestimated effect sizes and were rarely replicable in an independent sample.

We discuss the interpretation and implications of these findings within the context of accumulating evidence of the poor replicability of structural-brain-behaviour associations using grey matter volume, and more broadly of the replicability crisis of brain and behaviour sciences.

## Introduction

One striking fact when studying humans is the obvious inter-individual variability, be it in physical traits, such as hair, size, face,… or in psychological traits, such as personality, behaviour and cognition. This fascinating aspect of humankind has risen among scientists the quest for associations between different measurements. In particular, the early observations of inter-individual variability in human psychological skills and traits have triggered the search for correlates in brain structure, which can, for a few decades now, be measured in-vivo non-invasively. Relating interindividual variability in psychometric data to interindividual variability in brain structure is generally performed within the differential psychology perspective, that is, to better understand interindividual variability in behaviour. Nevertheless, structural-brain-behaviour (SBB) associations could also be of interest within a brain mapping perspective. From this later perspective the impact of the “found significant associations” usually remain within basic neuroscience theories of the relationships between brain organization and behavioural functions (Genon et al., 2018). In contrast, when performed within a differential psychology perspective, SBB findings often resonate beyond basic scientific literature as researchers infer about the “neural basis” of interindividual differences in traits and performance, such as personality or intelligence, which easily attract media and public attention. Thus, be it initially desired or not, the impact of SBB findings often goes beyond basic neuroscience and research communities.

Inter-individual variability in brain structure at the macroscale can be investigated in individual MRI anatomical scan with different types of measurements. The two most frequently used estimates of grey matter tissue’s features, with MRI technics, are grey matter volume and cortical thickness (Walhovd et al., 2017; Winkler et al., 2010). Both these markers have been used to identify associations between inter-individual variability in brain structure and interindividual variability in psychological outcomes. Grey matter volume estimation (GMV) (Good et al., 2001), as a relatively readily accessible method, has been a measure of choice for popular studies linking inter-individual variability of brain structure to individuals’ skills such as navigation expertise in London taxi drivers (Maguire, et al., 2006). In the same wave, inter-individual variability in GMV in different brain regions was also related to complex psychological constructs such a political orientation (Kanai et al., 2011) and social skills (“number of Facebook friends” (Kanai et al., 2012)). Hence, the old dream of differential psychology, finding its deepest roots into the phrenologist school of thoughts, was becoming true with the *scientific* evidence that “we are different because our brains are structurally different”.

However, a few years later, the replicability of some famous findings of association between GMV and psychological variables was scrutinized by (Boekel et al., 2015). The objective of the authors was to conduct a pure replication study of findings reported in the literature such as associations between the number of Facebook friends and GMV. Strikingly, their investigation revealed that most of the selected findings could not be replicated. At the same time, we explored associations between GMV and psychometric data from a different perspective (a brain mapping perspective), starting with a-priori defined regions of interest in the frontal lobe and searching for associations with a range of standard cognitive measures (Genon et al., 2017). This exploratory study mostly failed to show significant associations and when some were found, they could not be replicated in an independent sample. Subsequently, we performed a systematic and extensive evaluation of the replication rates of SBB associations using grey matter volume across a range of psychological measurements in a large sample of healthy adults which clearly confirmed our first exploratory findings: significant associations are very rare and when some can be found, despite with strict statistical control, they show very low replication rates (Kharabian Masouleh et al., 2019).

GMV is frequently seen in the neuroimaging community as an impure, multidetermined, and hence, crude estimate of grey matter tissue (Ashburner, 2009; Winkler et al., 2018). Since variabilities in the volumetric measures are mostly caused by surface area variations than by thickness variation itself, cortical thickness estimates (CT) may be seen as a more straightforward measure of brain structural features (Winkler et al., 2018, 2010). Accordingly, variability in cortical thickness could be expected to show relatively straightforward and reliable associations with behavioural measurements (But also see (Natu et al., 2019)). The scientific literature indeed provides a lot of apparent evidence of associations between local thickness and psychometric variables, in particular with measures of the big domains of differential psychology, namely intelligence(Choi et al., 2008; Karama et al., 2009; Menary et al., 2013; Schmitt et al., 2019) and personality (Hyatt et al., 2019; Owens et al., 2019; Riccelli et al., 2017a). Additionally, the scientific literature suggests that local variability in CT could be further related to more specific psychological aspects such as anxiety trait (Kühn et al., 2011), impulsivity (Schilling et al., 2012), musical perception performance (Foster and Zatorre, 2010) or mentalizing abilities (Rice and Redcay, 2015) just to name a few. Nevertheless, the question of the replicability of associations between psychometric data and local cortical thickness remains fully open. Evidencing replication rates of SBB with cortical thickness could clarify the broader question of the replication of SBB association in healthy adults.

In order to evaluate the replicability of associations between estimates of cortical thickness and behavioural measurements, in particular psychometric data, we performed an extensive empirical evaluation of the replicability rates of associations between common behavioural measurements and standard estimates of cortical thickness in a large, open-access high-quality datasets provided by the Human Connectom Project. Our evaluation was extensive at the behavioural level, including a wide range of psychological measures (i.e. psychometric variables). To systematically evaluate the replicability of SBB, we used two approaches for all SBB associations. First, for each behavioural variable, we searched for whole brain associations across 100 random samples of unrelated individuals and examined the spatial consistency of the findings of the cortical thickness-associations across the 100 samples. Second, for each behavioural variable we identified the clusters of vertices in the brain that were significantly associated with this variable in one of the previously mentioned random samples and then investigated whether we could replicate the found association, using a region of interest approach and several different criteria for replication, in an age- and sex-matched subsample. We also further investigated the influence of sample size on replicability. In line with our previous study using GMV(Kharabian Masouleh et al., 2019), for all approaches used here, we first provided the pattern of associations found with age, which can be considered as a reliably measured variable and reasonably expected to correlate with cortical thickness. Accordingly, thickness-associations with age, despite the narrow age-range in the human connectome project, serves here as a benchmark against which the replicability of SBB can be evaluated.

## Results

A total of 10200 exploratory whole brain SBB associations (each with 1000 permutations) were tested to empirically identify the replicability of the associations of 34 psychological scores with cortical thickness over 100 splits in independent matched subsamples, at three pre-defined sample sizes, within the HCP cohort (see Supplementary Table 1, for the total number of participants with available score for each of the psychological scores).

Altogether, in contrast to cortical thickness associations with age, significant SBB-associations were highly unlikely and only infrequently observed. For the majority of the tested psychological variables no significant association with cortical thickness were found in more than 90% of the whole brain analyses.

### SBB-associations among the healthy population

#### Replicability of “whole brain exploratory SBB-associations”

Ageing associated structural changes: Despite the limited age-range of participants within this sample (28-37 years), in line with previous studies (Fjell et al., 2014), vertex-wise associations of age with cortical thickness were widespread. For most subsamples, we found highly consistent negative associations of cortical thickness with age, in particular within the frontal lobe. Aggregate maps of spatial overlap of exploratory findings and density plots, summarizing the distribution of “frequency of significant findings” within each map are shown in figure 1A. When decreasing the sample size of the discovery cohort, the spatial overlap of significant age-associations over 100 splits decreased. More specifically, for the discovery sample of 294 participants, more than half of the significant vertices were consistently found as demonstrating significant association between cortical thickness and age in beyond 50% of the whole-brain exploratory analyses (i.e. rather high level of spatial consistency of significant findings). As the size of the subsamples decreased, the shape of the distribution also changed, and the median of the density plots fell around 30% and even 10% for samples consisting of 210 and 126 individuals, respectively.

**Figure 1.**
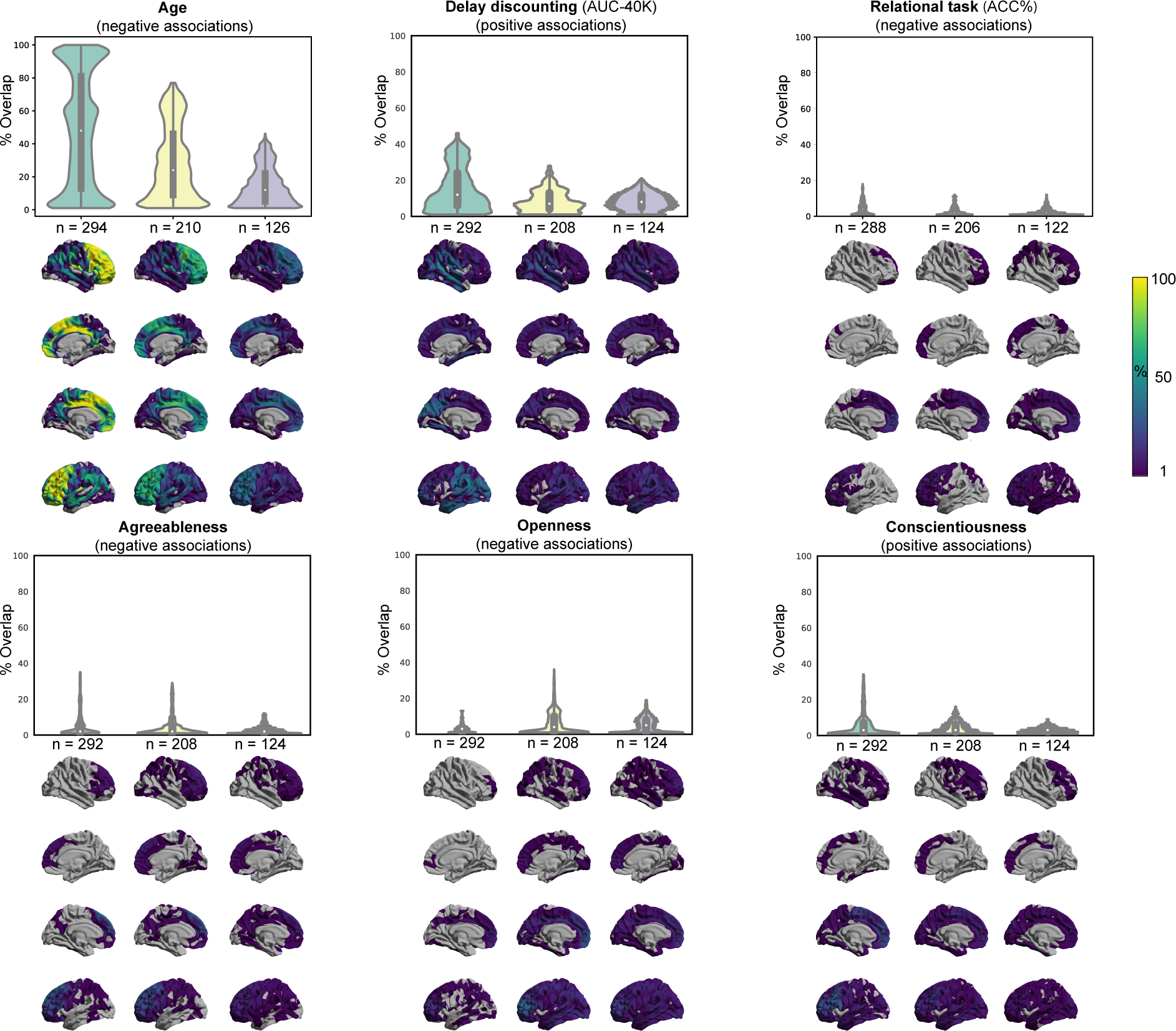
Replicability of exploratory results. Frequency of spatial overlap (density plots and aggregate maps) of significant findings from exploratory analysis over 100 random subsamples, calculated for three different sample sizes (x-axis). Here in addition to age, which is used as a benchmark, the top five behavioral scores with the highest frequency of overlapping findings are depicted. Brighter colors on spatial maps denote higher number of samples with a significant association at the respective vertex. AUC: Area under the curve; ACC: Accuracy.

These results highlight the influence of sample size on the replicability (frequency of overlap) of whole-brain significant associations, even for age, an objective measure that shows stable associations with variations in cortical thickness.

Structural associations of the psychological scores: In contrast, for most of the psychological scores, only few of the 100 discovery subsamples yielded significant clusters. Table 1 and supplementary Table 2 show the number of splits for which the exploratory whole-brain SBB-analysis resulted in *at least one* significant positively or negatively associated cluster for each score. These results reveal that finding significant SBB-associations using the exploratory approach in healthy individuals is highly *unlikely* for most of the psychological variables. Furthermore, the significant findings were spatially very diverse, that is, spatially overlapping findings were very rare.

**Table 1.**
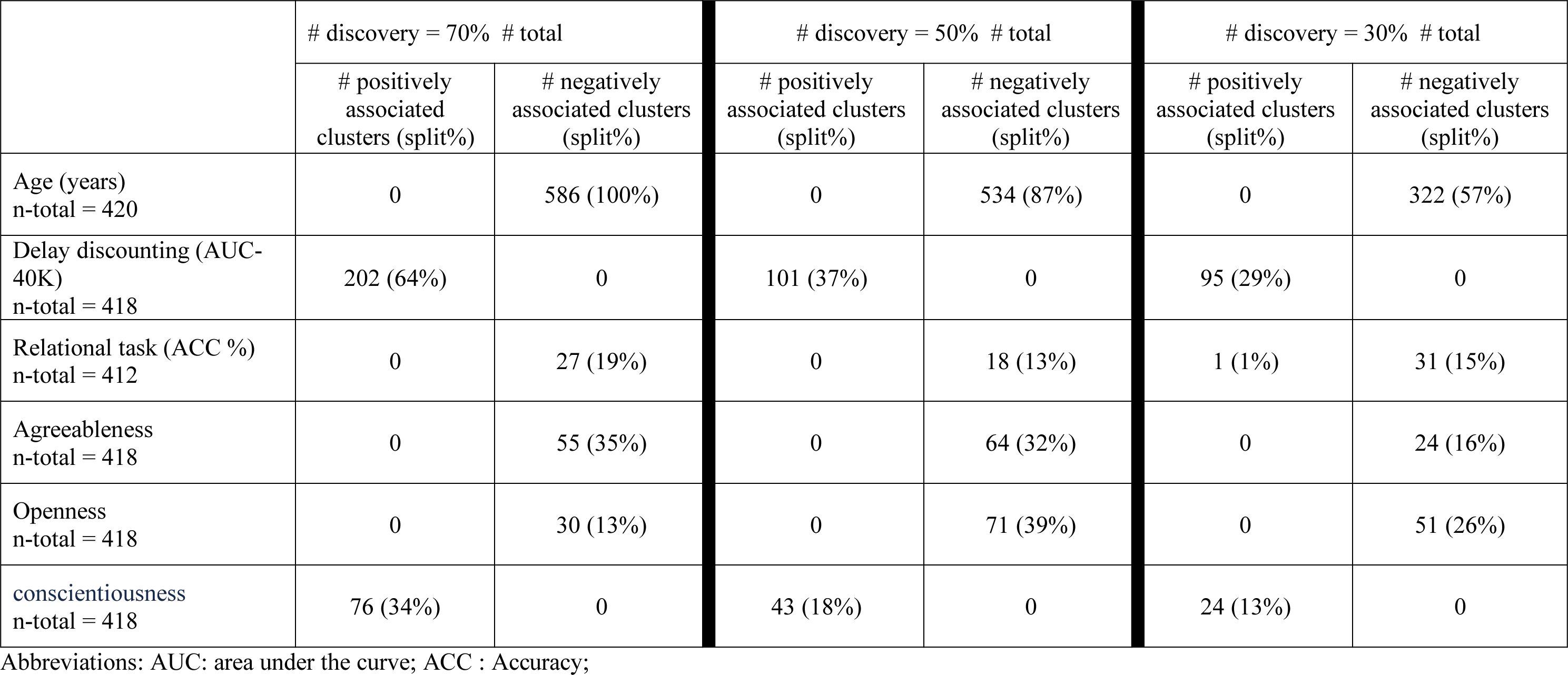
Summary of exploratory findings. For each discovery sample size, the number of clusters in which cortical thickness is positively or negatively associated with the tested phenotypic or psychological score is reported. The number of splits (out of 100) in which the clusters were detected are noted in parentheses (i.e. % of splits with at least one significant cluster [in the respective direction])

We here retained for further analyses the five psychological scores for which the discovery samples most frequently resulted in at least one significantly associated cluster. These three scores were the area under the curve for discounting of $40,000 (Delay discounting (AUC-40K)), the accuracy percentage during the relational blocks from the in scanner relational task (Relational task (accuracy percentage)), and agreeableness, openness and conscientiousness scores of the five factor personality model. For example, for the discovery samples of 292 adults, in 64 out of 100 randomly generated discovery samples, at least one cluster (not necessarily overlapping) showed a significant positive association between area under the curve for discounting of $40K and cortical thickness (Table 1)).

Yet again, in line with our observations for age associations, generally, the probability of finding at least one significant cluster tend to decrease in smaller discovery samples (see Table 1). Likewise, as the discovery sample size decreased, the maximum rate of spatial overlap, as denoted by the height of the density plots, decreased (see Figure 1B-F). The width of these plots show that the majority (> 50%) of the significant vertices spatially overlapped only in less than 10% of the discovery samples. In the same line, the variability depicted by the spatial maps highlight that many vertices are found as significant only in one out of 100 analyses.

These results highlight that finding a significant association between normal variations on psychometric data and vertex-wise measures of cortical thickness among healthy individuals is highly unlikely, for most of the tested domains. Furthermore, they underscore the extent of spatial inconsistency and the *poor replicability* of the significant SBB-associations from *exploratory analyses*.

#### Confirmatory ROI-based SBB-replicability

Age effects: Over all three tested sample sizes, in more than 99% of the a-priori defined ROIs, age associations were found to be in the same “direction” in the discovery and test samples (i.e. replicated based on “sign” criteria). The examination of replicated findings based on “statistical significance” revealed replicated effects in more than 84% of ROIs. This rate of ROI-based replicability increased from ~84% to 91%, as the test sample size increased from 126 to 294 individuals (see figure 2). Furthermore, as the dark blue segments in the outer layers of figure 2 indicate, Bayesian hypothesis testing revealed moderate-to-strong evidence for H1 in more than 63% of the ROIs.

**Figure 2.**
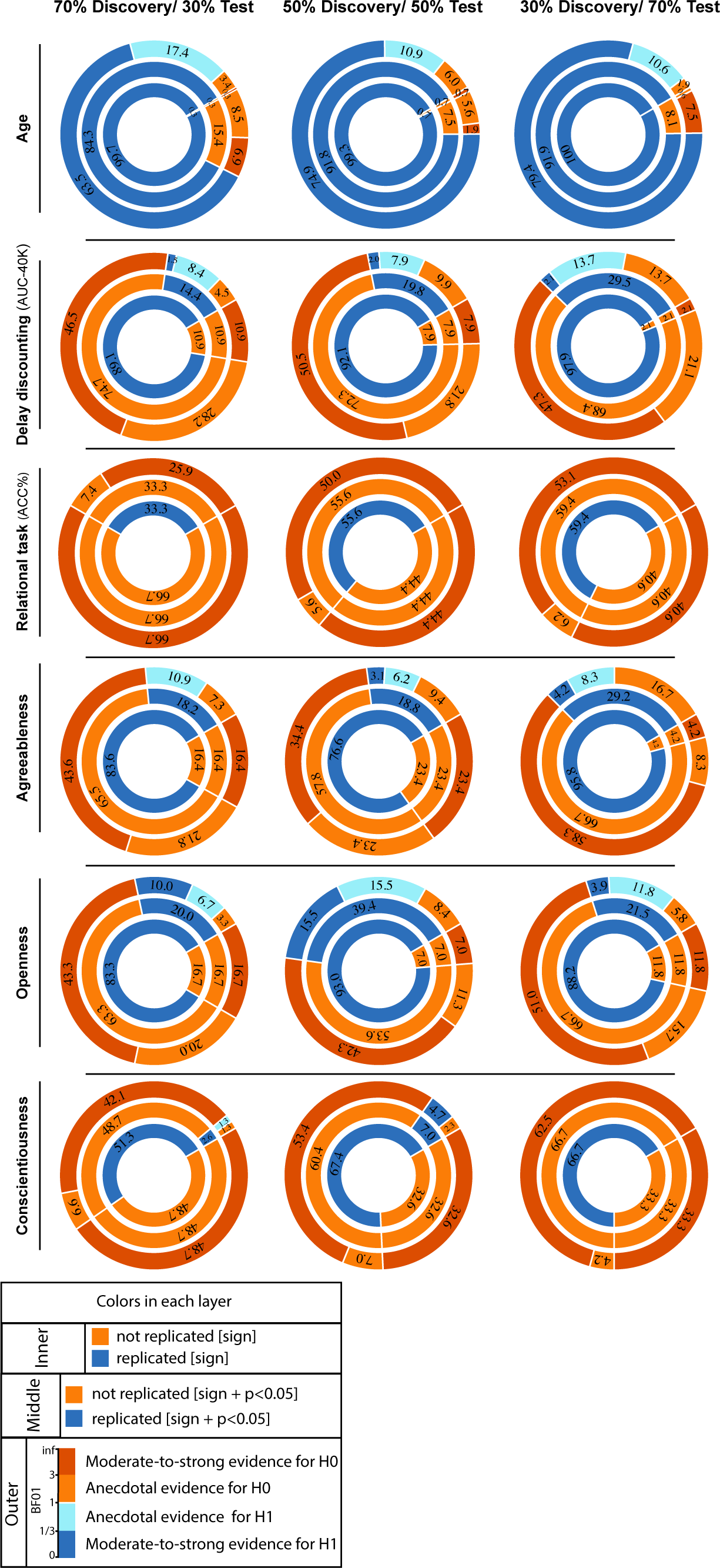
ROI-based confirmatory replication results. Donut plots summerising ROI-based replication rates (% of ROI) using three different critera for three different sample sizes among heathy participants. The most inner layers depict replication using “sign” only (blue: replicated, orange: not replciated). The middle layers define replication based on similar “sign” as well as “statistical significance” (i.e. p < 0.05) (blue: replicated, orange: not replciate). The most outer layers define replication using “bayes factor” (blue: “moderate-to-string evidece for H1, light blue: anecdotal evidence for H1; light orange: anecdotal evidence for H0, orange: “moderate-to-string evidece for H0);

Psychological variables: Figure 2 also illustrates the replicability rates of structural associations of the top five psychological measures from the whole brain analyses (the area under the curve for discounting of $40K, accuracy percentage of the relational task, and the three personality scores: agreeableness, openness and conscientiousness).

Despite the mean thickness associations of delay discounting (AUC-40K) being in the same direction in the discovery and test samples (positive SBB-association), for the majority of the ROIs (>89%), only less than 30% of all ROIs showed replicated effects based on “statistical significance” criterion. Finally, less than 3% of the ROIs were identified as “successfully replicated” based on the Bayes factors. (Figure 2).

For the three tested samples sizes, associations of the accuracy percentage of the relational task and cortical thickness were in the same direction (positive SBB-association) in the discovery and test pairs among beyond 33% of ROIs. Nevertheless, associations within none of these ROIs were defined as successfully replicated using the statistical significance or the Bayes factor criteria (Figure 2).

Among the three top associated personality scores, negative associations between agreeableness and cortical thickness were in the same direction as the discovery effects among more than ~77% of ROIs. The significant-replication was found among 18% to 30% of all ROIs, for the three tested sample sizes, and Bayes factors defined a successful replication in less than 5% of all ROIs. Negative correlations between openness scores and average cortical thickness were depicted in ~90 % of the ROIs, but significant-replication was found in 20% to 40% of all ROIs, for the three test sample sizes. Along the same line, successful replication based on the Bayes factors was below 16%. Finally, paired-test samples confirmed direction of association between conscientiousness and cortical thickness in less than 52% of ROIs. Significant replication was found within not more than 7% of ROIs and less than 5% were defined as successfully replicated in the replication sample using the Bayes factor criteria (Figure 2).

In general, these results show the span of replicability of structural (thickness) associations from highly replicable age-effects to very poorly replicable psychological associations, within the HCP cohort, consisting of young, healthy adults. They also highlight the influence of the sample size, as well as the criteria that is used to define successful replication on the rate of replicability of SBB-effects in independent samples.

#### Effect size in the discovery sample and its link with effect size of the test sample and actual replication

Figure 3 plots discovery versus replication effect size (i.e. correlation coefficient) for each ROI and for three test sample sizes. Focusing on by-“sign” replicated ROIs (blue), for the five psychological scores (delay discounting (AUC-40K), relational task (relative accuracy), and the three personality scores: agreeableness, openness, conscientiousness) revealed that the discovery samples resulted in overall larger effects (magnitude of “original effects”) compared to the test samples (Replication effects). Indeed, the marginal distributions are centred around smaller correlation coefficients in the y-dimension (test sample) compared to the x-axis (discovery samples). Furthermore, for these by-“sign” replicated ROIs (blue dots), positive relationship between the effect sizes of the behavioural associations in the discovery and test samples occurred rarely (blue lines in each subplot).

**Figure 3.**
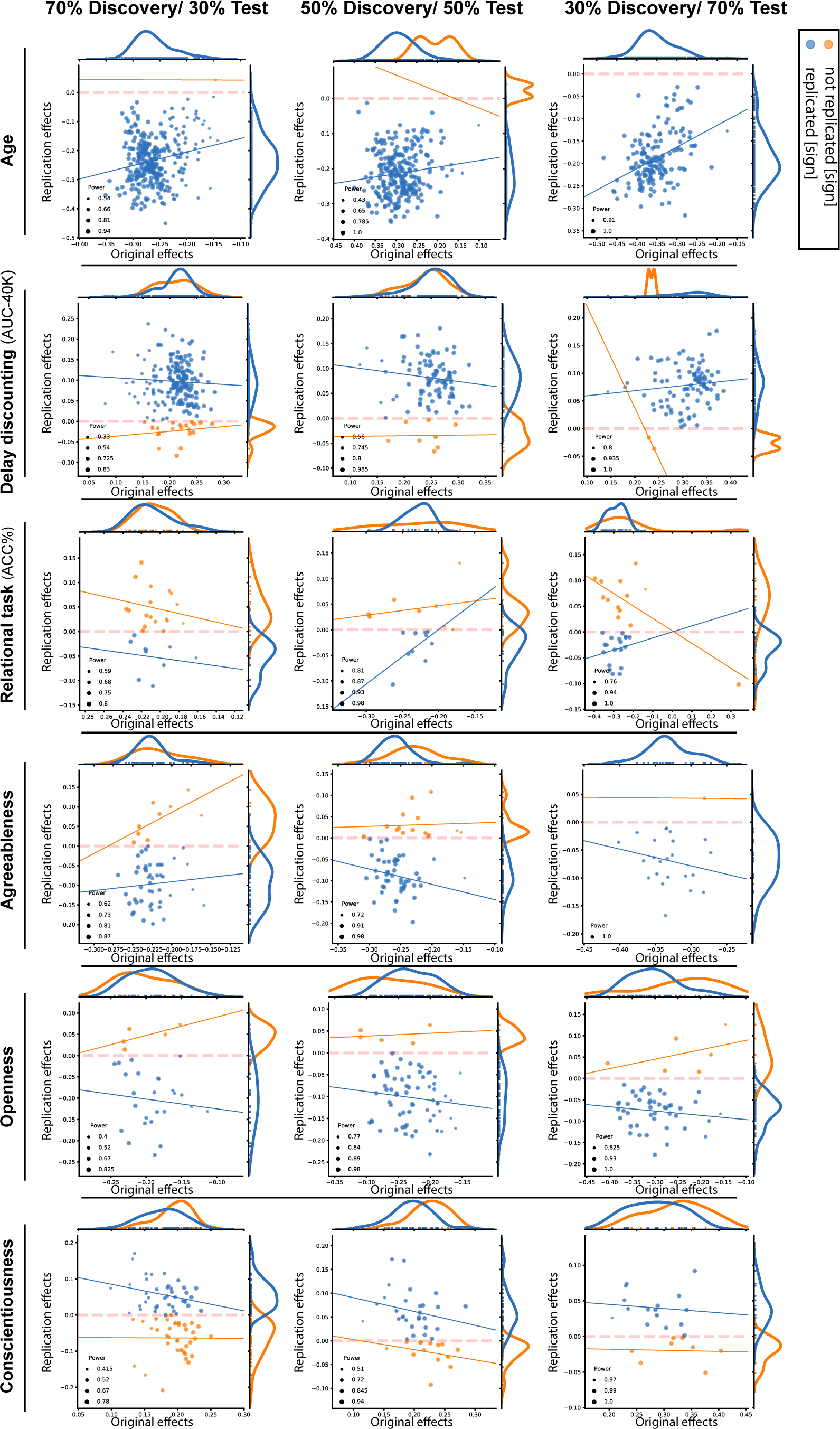
Discovery versus replication effects sizes. Scatter plots of effect sizes in the discovery versus replication sample for all ROIs from 100 splits within healthy cohort; each point denote one ROI, which is color-coded based on its replciation status (by-“sign”). Size of each point is proportional to its estimated statistical power of replication. Regresion lines are drawn for the replciated and unreplicated ROIs, separately.

In contrast, for age, the effect sizes of the discovery and test pairs were generally positively correlated, suggesting that the regions with greater negative structural association with age in the discovery sample, also tended to show stronger negative associations within the matched test sample.

To investigate if the replication power, estimated using the correlation coefficient within the discovery samples, was linked to a higher probability of *actual* replication in the test samples, the ROIs were grouped into replicated and not-replicated, based on the “statistical significance” criterion (p < 0.05). While for age associations the estimations of statistical power were generally higher among the replicated compared to not-replicated ROIs (p-value of the Mann-Whitney U tests < 10^−2^), for structural associations of the psychological scores, this was not the case. These findings highlight the unreliable aspect of effect size estimations of SBB-associations within the discovery samples among healthy individuals, which further result in uninformative estimated statistical power.

## Discussion

Overall our empirical evaluation of the replicability of associations between *in-vivo* estimates of cortical thickness and psychometric variables in healthy adults confirmed the results of our previous examination using neuroimaging estimates of grey matter volume (Kharabian Masouleh et al., 2019). It should be noted that the current conceptual replication of our previous findings has been performed in a different cohort of healthy adults than our previous study, here capitalizing on acknowledged high-quality neuroimaging data. Despite this optimization of our experimental setting, our extensive evaluation across a range of psychometric variables reveals that significant SBB associations using *in-vivo* measurements of cortical thickness are very unlikely. When significant associations were found, these associations show very poor replication rates. In line with our previous study, performed using grey matter volume, we here also highlighted the influence of sample size on the replication rates. Hence, overall, the current study extends our previous alarming findings on the poor replicability of SBB studies to experimental design using cortical thickness estimates. Below we discuss the interpretation and implications of these findings, especially in the context of the replicability crisis of psychological, social and neuroimaging-based neurosciences. Based on this discussion and acknowledging the potential contribution of SBB associations studies to our understanding of brain-behaviour relationships, we finally propose some possible recommendations for individual studies.

### Unlikely associations between local cortical thickness and psychological variables

In this study, we searched for significant associations between inter-individual variability in cortical thickness across the cerebral cortex surface and inter-individual variability in 34 psychometric variables. For most of the examined psychometric variable (27 out of 34), significant associations were found in less than 10% of our exploratory analyses. Even though the sensitivity of the psychological measures to detect relevant variations in cognition and personality could be questioned, the reliability and relative validity of these tests have been carefully ensured by the HCP initiative (Barch et al., 2013) and, accordingly, could not account for the poor rates of significant SBB findings. A relatively low variability due to a potential selection bias of the participants (subsamples from 420 unrelated individuals) could partly explained the high rates of null findings, nevertheless our results also show that, despite the age range is relatively restricted, it shows highly significant replicable associations with CT. Thus, the data here reveals that, although there is inter-individual variability in psychometric data and estimates of cortical thickness, significant associations between specific brain regions and specific psychometric variable can hardly be evidenced. However, several scientific publications report significant relationships between behavioural variables (Foster and Zatorre, 2010; Kühn et al., 2011; Rice and Redcay, 2015; Schilling et al., 2012), in particular psychometric intelligence-related (Karama et al., 2011; Schnack et al., 2015) and personality-related (Owens et al., 2019; Riccelli et al., 2017b) measures and local thickness. Considering that in the current state-of-the-art, visibility is given mainly to studies which can report a significant finding (Dwan et al., 2013; Franco et al., 2014; Nissen et al., 2016), leaving the potentially highly numerous null findings completely unknown, our in-depth investigations suggests that the current picture of associations between thickness of the cortex and psychometric variables in the scientific and general collective mind could be highly flawed.

### Spatially inconsistent patterns across the brain with an exploratory approach

In a typical exploratory approach, one or several given psychometric variables of interest are a-priori defined and a mass-univariate analysis, in which statistical associations is tested for each vertex or each parcel of the brain, is performed. The regions that survive statistical control for multiple testing are then reported. Importantly, traditionally, SBB studies searching for associations between psychometric variables and cortical thickness use low-resolution brain atlases that reflects mainly macro-anatomical boundaries (such as Desikan-Killiany adult cortical atlas (Desikan et al., 2006), dividing the whole cortex into 68 regions), instead of voxel/vertex-wise data or an atlas based on functional subdivision (for a review see (Eickhoff et al., 2018)). This common approach, in which a broad summarization into macroanatomical territories is imposed to the data, may in turn increase the likelihood of finding irreproducible spatial associations.

Nevertheless, until recently, replication of such studies and the evaluation of reproducibility of the found associations received little attention. In the current study, we chose a high resolution, vertex-wise approach and by using random splits of a large cohort, we could demonstrate that for a given psychometric variable the spatial pattern of associations usually varies across the generated random samples. Overall, the maximum overlap of significant associations of psychometric measures, across 100 splits, was less than 50% and for most brain regions was much lower. Considering one of the standard personality factors, for example agreeableness, this suggests that if 100 studies, with an identical experimental settings using a mass-univariate approach and within the same population, search for its associations with cortical thickness, an association with cortical thickness in the a given region (such as the temporoparietal junction that was found here) would be reported only in five studies. These findings suggest that the associations between brain regions and specific psychological measurements should be taken with extreme caution and inferences about associations between brain structures and behavioural variables, in particular in personality traits should be seriously questioned.

### Region-of-interest-based replication attempts in paired samples predominantly failed

Estimates of brain structure, such as grey matter volume and cortical thickness are known to vary with demographical variables such as age and gender (Fjell et al., 2014; Savic and Arver, 2014). Accordingly, it could be argued that the poor consistency of the spatial pattern of cortical thickness associations with psychological variables across random splits could be due to the diversity of the splits with regards to these variables. Furthermore, it could be argued that the common corrections for multiple comparison, while importantly limiting the chance of false positive findings (Eklund et al., 2016), might be too stringent and thus results in increased false negative rates (Cremers et al., 2017; Noble et al., 2019) and decreased probability of overlapping findings within the exploratory analyses. To address these two issues, in this study, we also examined replications of associations using a *confirmatory* approach, within demographically paired samples. Concretely, when associations were found between cortical thickness and psychometric variables with the exploratory approach, the associations between thickness in the found brain clusters and the psychometric variable were subsequently tested in a demographically matched sample. In line with the spatial consistency investigation, this confirmatory approach showed that the significant associations could not be replicated in the vast majority of the identified clusters and that the data supported evidence in favour of the null hypothesis (of no association) in the matched samples.

Thus, altogether, our study suggests that significant associations between local cortical thickness and psychometric variables are relatively unlikely to be found and when some significant associations are found, they are predominantly not replicable even within an identical experimental setting. Furthermore, our observation of a lack of clear positive association between the original and replication effect size within by-sign replicated clusters as well as our observation of no clear association between the replication power and the rate of significantly replicated clusters, further suggest the influence of random noise. This pattern appears particularly in the identification of associations within small samples, where it results in spurious findings and overestimation of effect size and hence unreliable estimates of replication power (Albers and Lakens, 2018). Hence the patterns of findings suggests that, when an association was found, it might have been driven by random noise (Loken and Gelman, 2017) (and winner’s curse (Sham and Purcell, 2014)).

### A replication crisis for structural brain-behaviour studies ?

To summarize the current state of the art regarding the replicability of SBB studies, In 2015, Boekel and collaborators demonstrated that among a few selected structural associations from the previously published studies, most of them could not be replicated. In our previous studies (Genon et al., 2017; Kharabian Masouleh et al., 2019) and the current one, we extended the scope of these findings. We showed that, whatever the neuroimaging measure of brain structure used, be it, grey matter volume or cortical thickness 1) the rate of significant associations between local brain structure and psychometric scores is extremely low; 2) the rate of replication of the found significant associations (after strict statistical threshold), using both exploratory and confirmatory approach is extremely low; 3) replication rate decreases as sample size decreases. Importantly, in our previous study, as well as the current one, we had several quality check and associations between age and brain structure served as a benchmark. These control conditions support the validity of the neuroimaging measures and the statistical approach used in our analyses. It is worth noting as well that our studies showing the low replicability of SBB using voxel-based morphometry and the low replicability of SBB using cortical thickness estimates were performed in two different datasets. In other words, the poor replication rates of SBB can hardly be attributed to a questionable quality of the datasets, neither to a questionable validity of the brain structural and behavioural variables. In our previous study, we also demonstrated that the same approach used within a clinical population yielded higher replicability rates suggesting that structural variability (atrophy) and symptomatic behaviour can be reliability related to each other (Kharabian Masouleh et al., 2019). We therefore here conclude that the replicability of the standard SBB in healthy populations, and hence the validity of this approach in a differential psychology perspective, should be finally questioned.

To understand the impact of our findings, it should be kept in mind that we here refer specifically to univariate analysis linking local brain structural measurements to psychometric variables in healthy adult samples and leading to conclusions in the vein of “bigger brain volume in region X, is associated to psychological trait Y or cognitive performance Z”. These types of inferences are mainly elaborated within a differential psychology framework and it is generally from this framework that they propagate into the collective mind. Thus, the associations revealed by standard SBB studies in healthy adults should be questioned and care should be taken that inferences based on not-yet replicated findings do not propagate beyond the scientific community.

### Conclusions, recommendations and available resources

In the previous years, the replicability crises that has shaken social (Camerer et al., 2018; Vul et al., 2009), psychological (Franco et al., 2014; Lindsay, 2015; Open Science Collaboration, 2015), biomedical and neuroimaging-based neuroscience (Button et al., 2013; Eklund et al., 2016; Grabitz et al., 2018; Poldrack et al., 2017) have contributed to the establishment of several recommendations or guidelines, e.g. (Button et al., 2013; Munafò et al., 2017; Poldrack et al., 2017; Wagenmakers et al., 2012). Many of these recommendations are directly relevant in the context of SBB. First, at dataset level, considering the central role of sample size and related power issues in the replication crisis (Kharabian Masouleh et al., 2019; Noble et al., 2019), in line with evidence provided in our current study, identification of robust links between psychological variables and brain phenotype needs moving towards big data samples, e.g. (Miller et al., 2016; Van Essen et al., 2013). Second, because false positive appear to occur more frequently and be unnoticed when the degrees of freedom and the exact analyses path leading to the published findings are undisclosed (Button et al., 2013; Gelman and Loken, 2014), analysis should be justified and documented and disclosure should be increased. Along the same line, the exploratory nature of the analysis should be acknowledged. In particular, considering that the SBB approach is typically an observational empirical approach, it is frequent that the data has not been acquired for the purpose of the finally reported SBB analysis.

In this case the exploratory nature of the investigation should be clearly acknowledged. However, if the study was truly a confirmatory study, it should be pre-registered (Chambers et al., 2015; Nosek et al., 2015). The Open Science Framework (OSF, (Foster, MSLS and Deardorff, MLIS, 2017)) and many scientific journals offer registration frameworks for empirical research. Furthermore, ideally, the traceability and availability of data and analyses should be ensured. Related to the degree of freedom in the analyses pipelines and computation tools, direct access to the code and data should be provided. SBB analyses being often based on correlational approaches, they are for example highly susceptible to the effect of outliers. Ideally the reader should be allowed to directly visualize the findings for his/her own insight (such as on scatter plots) and, additionally, to evaluate how the results are affected, for example, by a few subjects exclusion. Freely available resources such as knitr (https://yihui.org/knitr/) and datalad (https://www.datalad.org) can be used to promote transparency and traceability and data sharing platforms such as OpenNEURO (https://openneuro.org) and Neurovault (Gorgolewski et al., 2015) offers free access to data and results of statistical maps. Finally, for any found significant association, replication studies in an independent dataset should be performed. We note however, that the concretization of this suggested good research practice is highly dependent on the active contributions of publishers and funding organizations by providing incentives for replications effort. While these practices could be seen as time consuming and resource consuming in the short-term, they will all contribute to a better understanding of brain-behaviour relationships in the future, but also more generally contribute to build a more optimistic, fruitful and ethical scientific culture in the long run.

## Methods

### Participants

Healthy adults’ data were selected from the publicly available data from the Human Connectome Project (HCP; http://www.humanconnectome.org/), which comprised data from 1206 individuals (656 females), 298 MZ twins, 188 DZ twins, and 720 singletons, with mean age 28.8 years (SD = 3.7, range = 22–37). After passing the HCP quality control and assurance standards (Marcus et al., 2011), structural data of 1113 individuals were released. The full set of inclusion and exclusion criteria are described elsewhere (Glasser et al., 2013; Van Essen et al., 2013). Here we selected a subset of unrelated individuals from this cohort, consisting of 420 individuals (age: 28 ± 3.7, 210 female), with good quality structural scans.

### Phenotypical measurements

#### Non-psychological measurements

Age is a reliably measured variable, whose impact on the neuroimaging derived markers of micro- and macrostructure of the brain have been rigorously studied and robustly demonstrated within different cohorts of healthy individuals (Campbell et al., 2015; Fjell et al., 2014; Raz, 2005; Walhovd et al., 2017). Therefore, in line with our previous study (Kharabian Masouleh et al., 2019), we here used age-associations with the estimates of the cortical thickness as a benchmark against which we compare the replicability of behavioural associations.

#### Psychological measurements

The psychological measurements consisted of a subset of 34 standard psychometrics and neuropsychological tests, available in the HCP cohort. The testing consisted of the following main categories:

##### Emotion

Emotion recognition from the Penn emotion recognition test battery (Gur et al., 2010). Negative affect (anger), psychological well-being (life satisfaction) and self-reported measure of emotional support acquired using the NIH toolbox surveys.

##### Cognition

Card sorting test from the NIH toolbox, measuring executive cognitive flexibility. Area under the curve for discounting of $200 and $40,000 (Estle et al., 2006; Myerson et al., 2001), as summary measures assessing self-regulation and impulsive behaviour. Reaction time and total number of correct responses from the Penn word memory test, measuring verbal episodic memory. List sorting from NIH toolbox, measuring working memory. Number of correct responses in Penn progressive matrices, measuring fluid intelligence.

Also, few composite cognitive scores were created by averaging various cognitive tests: Crystallized composite score, which is assessed by averaging the normalized scores of each of the NIH Toolbox tests that are crystallized measures (Picture Vocabulary and Reading Tests), measuring verbal reasoning. Early childhood composite cognitive score, which is assessed by averaging the normalized scores of the cognitive measures that comprise the Early Childhood Battery (Picture Vocabulary, Flanker, Dimensional change card sorting and Picture Sequence Memory). The early childhood composite score provides a reliable overall snapshot of general cognitive functioning. Finally, by averaging the normalized scores of each of the fluid and crystallized cognition measures, a total composite score of cognition is generated, measuring levels of cognitive functioning.

In addition to above-mentioned measures, performance (average accuracies and median reaction times) of few in-scanner tasks acquired during functional MRI sessions, were used as additional cognitive scores, consisting of working memory task (two-back working memory tasks for faces, body, tools and places), language task (math) as well as average accuracy of the relational processing task (Smith et al., 2007).

##### Personality

Personality was assessed using five factor personality model of personality (Gur et al., 2010).

Furthermore, handedness and dexterity were added as basic motor functions measurements. Supplementary Table 1 demonstrates information about the distribution of the selected scores within the sample of 420 HCP individuals.

### MRI acquisition and preprocessing

Two high-resolution (isotropic 700 µm) 3D MPRAGE T1-weighted images were acquired using the HCP’s custom 3T Siemens Skyra scanner with the following parameters: TE/TR/TI = 2.14/2400/1000 ms, field of view (FOV) =224 mm, flip angle=8°, Bandwidth (BW)=210 Hz per pixel. Two T2-weighted images were also acquired with identical geometry (TE/TR = 565/3200 ms, variable flip angle, BW = 744 Hz per pixel). Full description of MRI protocols of the HCP is previously described in (Glasser et al., 2013). Images underwent gradient nonlinearity correction (using acquired B1 bias field) and the two scans of each modality were co-registered and averaged. Cortical thickness estimates derived using FreeSurfer v5.3-HCP pipeline (https://github.com/Washington-University/HCPpipelines/blob/master/FreeSurfer/FreeSurferPipeline-v5.3.0-HCP.sh), were downloaded from the Amazon Web Services. The outcome of this pipeline (Glasser et al., 2013), which is optimized for the HCP data and further incorporates T2-weighted images into the FreeSurfer analysis pipeline, is frequently used to report cortical thickness variations within the HCP sample.

To perform group level analysis, each individual’s thickness estimations were registered to the fsaverage surface (fsaverage - 163,842 vertices per hemisphere), through a non-linear surface-based inter-subject registration procedure that aligns the cortical folding patterns of each subject to a standard surface (fsaverage) space (Fischl et al., 1999).

Finally each individual’s thickness estimates were mapped to the fsaverage surface and were spatially smoothed on the surface using a gaussian kernel of 15 mm (full-width-half-maximum).

### Statistical analysis

SBB-associations are commonly derived in an exploratory setting using a mass-univariate approach, in which a linear model is used to fit interindividual variability in the psychological score to measure of brain structure, here cortical thickness, at each vertex. Inference is then usually made at cluster level, in which groups of adjacent vertices that support the link between cortical thickness and the tested score are clustered together.

Replicability of thus-defined associations could be assessed by conducting a similar whole-brain vertex-wise exploratory analysis in another sample of individuals and comparing the spatial location of the significant findings that survive multiple comparison correction, between the two samples (e.g. (Martínez et al., 2015)). Alternatively, replicability could be assessed, using a confirmatory approach, in which only regions showing significant SBB-association in the initial exploratory analysis, i.e. regions of interest (ROIs), are considered for testing the existence of the association between brain structure and the same psychological score in an independent sample (e.g. (Boekel et al., 2015)). The latter procedure commonly focuses on a summary measure of structure within each ROI and tests for existence of the SBB-association in the direction suggested by the initial exploratory analysis. Thus this approach circumvents the need for multiple comparison correction and therefore increases the power of replication. In line with our previous study (Kharabian Masouleh et al., 2019), here we assessed replicability of associations between each behavioural measure and grey mater structural variability, using both approaches: the whole brain replication approach and the ROI replication approach, which are explained in details in the following sections.

#### Replicability of whole brain exploratory SBB-associations

Whole-brain GLM analyses: 100 random subsamples (of same size) were drawn from the main cohort. Hereafter, each of these subsamples is called a “discovery sample”. In each of these samples, SBB-associations were identified using the vertex-wise exploratory approach after controlling for confounders. This was done by using the general linear model (GLM) as implemented in the “PALM” tool (https://fsl.fmrib.ox.ac.uk/fsl/fslwiki/PALM), with 1000 permutations. Age, sex and education were modelled as confounders.

Inference was then made using threshold-free cluster enhancement (TFCE) (Smith and Nichols, 2009), which unlike other cluster-based thresholding approaches, does not require an arbitrary a-priori cluster forming threshold. Significance was set at P < 0.05, corrected for two sided nature of the tests (positive and negative associations) as well as the two hemispheres.

Spatial consistency maps and density plots: To demonstrate the spatial overlap of significant associations over 100 subsamples, spatial consistency maps were generated. To do so, the binarized maps of all clusters on the surface that showed significant association in the same direction between each psychological score and cortical thickness were generated (i.e. vertices belonging to a significant cluster get the value “1” and all other voxels were labelled “0”) and added over all 100 subsamples. These aggregate surfaces denote the frequency of finding a *significant* association between the behavioural score and cortical thickness, at each vertex. Accordingly, a vertex with value of 10 in the aggregate surface map has been found to be significantly associated with the phenotypical score in 10 out of 100 subsamples. Density plots were also generated to represent the distribution of values within each such map, i.e. the distribution of “frequency of significant finding”. Hence, the spatial vertex-wise “significance overlap surface maps” as well as density plots of the distribution of values within each map give indications of the replicability of “whole brain exploratory SBB-associations” for each psychological score.

#### Replicability of SBB-associations using confirmatory ROI-based approach

ROI-based confirmatory analyses: The replicability of the SBB associations was also evaluated with the ROI-based confirmatory approach. For each of the 100 discovery subsamples, an age- and sex-matched “test sample” was generated from the remaining participants of the main cohort.

In this analysis, for each psychological variable, the significant clusters from the above-mentioned exploratory approach from every “discovery sample” were used as a-priori ROIs. Average cortical thickness over all vertices within the ROI was then calculated for each participant in the respective “discovery” and “test” pair subsamples. Within each subsample, association between the average cortical thickness and the psychological variable was assessed using ranked-partial correlation, controlling for confounding factors. The correlation coefficient was then compared between each discovery and test pair, providing means to assess “ROI-based SBB replicability” rates for each psychological score. Accordingly, each ROI was examined only once, to identify if associations between average thickness in this ROI and the psychological score from the discovery subsample could be confirmed in the paired test sample. Replicability rates were quantified according to different indexes (see below) over all ROIs from 100 discovery samples, yielding a percentage of “successfully replicated” surface ROIs based on each index.

Indexes of replicability:

##### Sign

This lenient definition of replication compares only the sign of correlation coefficients of associations within each ROI between the discovery and the matched-test sample. Accordingly, any effect that was in the same direction in both samples (even if very close to zero) is defined as a “successful” replication using this criterion.

##### Statistical Significance

Another straightforward method for evaluating replication simply defines statistically significant effects (e.g. p-value < 0.05) that are in the same direction as the original effects (from the discovery sample) as “successful” replication. This criteria is consistent with what is commonly used in the psychological sciences to decide whether a replication attempt “worked” (Open Science Collaboration, 2015). Yet, a key weakness of this approach is that it treats the threshold (p < 0.05) as a bright-line criterion between replication success and failure. Furthermore, it does not quantify the decisiveness of the evidence that the data provides for and against the presence of the correlation (Boekel et al., 2015; Wagenmakers et al., 2015). However, such an estimation can be provided by using the “Bayes factors”.

##### Bayes Factor

To compare the evidence that the “test subsample” provided for or against the presence of an association (H1 and H0, respectively), we additionally quantified SBB-replication within each ROI, using Bayes factors (Jeffreys, 1961). Similar to Boekel et al. (2015), here we used the adjusted (one-sided) Jeffry’s test (Jeffreys, 1961) based on a uniform prior distribution for the correlation coefficient. As we intended to confirm the SBB-associations defined in the discovery subsamples, the alternative hypothesis (H1) in this study was considered one-sided (in line with Boekel et al. (2015)).

In line with our previous study (Kharabian Masouleh et al., 2019), Bayes factors (BF) were summarized into four categories as illustrated in the bar legend of Figure 2. These categories are used to facilitate the interpretation and comparison of replication rates. For example, a BF_01_ lower than 1/3 shows that the data is three times or more likely to have happened under H1 than H0. “Successful” replication is defined as a BF_01_ lower than 1/3.

#### Investigation on factors influencing replicability of SBB-associations among healthy individuals

Sample size: In order to study the influence of sample size on the replicability of SBB-associations, for each psychological measure, the main sample was divided into discovery and test pairs at three different ratios: 70% discovery and 30% test, 50% discovery and 50% test and finally 30% discovery and 70% test. As mentioned earlier, in each case, the discovery and test counterparts were randomly generated 100 times in order to quantify the replication rates. For example, to assess the replicability of brain structural associations of age, in the case of “70% discovery and 30% test”, the entire HCP sample (n = 420, unrelated individuals) was divided into a discovery group of n = 294 participants and an age- and sex-matched test pair sample of n = 126 and this split procedure was repeated 100 times. Similarly, for generating equal-sized discovery and test subsamples, 100 randomly generated age and sex matched split-half samples were generated from the main cohort.

Effect size: Furthermore, to study the influence of the effect size on the replication rates, we focused on the effect sizes within each a-priori ROI in the discovery samples. Here we tested the following two assumptions:

1) ROIs with larger effect sizes in the discovery sample result in larger effect sizes in the test sample pairs (i.e. positive association between effect size in the discovery and test samples).

2) ROIs with larger effect sizes in the discovery sample are more likely to result in a “significant” replication in the independent sample.

To test the first assumption, in the “ROI-based SBB-replicability” the association between effect size in the discovery and test pairs were calculated for each psychological measure. These associations were calculated separately for the replicated (defined using “sign” criterion) and not-replicated ROIs. We expected to find a positive association between discovery and confirmatory effect sizes, for the “successfully replicated effects”.

To test the second assumption, for each ROI, we calculated its replication statistical power and compared it between replicated and not-replicated ROIs (here replication was defined using “Statistical Significance” criterion). The statistical power of a test is the probability that it will correctly reject the null hypothesis when the null is false. In a bias-free case, the power of the replication is a function of the replication sample size, real size of the effect and the nominal type I error rate (α). In this study, the replication power was estimated based on the size of the effects as they were defined in the discovery sample and a significant threshold of 0.05 (one-sided) and was calculated using “pwr” library in R (https://www.r-project.org).

These analyses were performed for each discovery-test split size, separately (i.e. 70%-30%, 50%-50% and 30%-70% discovery-test sample sizes, respectively).

## Acknowledgments

This work was supported by the Deutsche Forschungsgemeinschaft (DFG, GE 2835/1-1, EI 816/4-1), the Helmholtz Portfolio Theme ‘Supercomputing and Modelling for the Human Brain’ and the European Union’s Horizon 2020 Research and Innovation Programme under Grant Agreement No. 785907 (HBP SGA2). Data were provided by the Human Connectome Project, WU-Minn Consortium (Principal Investigators: David Van Essen and Kamil Ugurbil; 1U54MH091657) funded by the 16 NIH Institutes and Centers that support the NIH Blueprint for Neuroscience Research; and by the McDonnell Center for Systems Neuroscience at Washington University.

## References

Albers C, Lakens D. 2018. When power analyses based on pilot data are biased: Inaccurate effect size estimators and follow-up bias. J Exp Soc Psychol 74:187–195. doi:10.1016/j.jesp.2017.09.004

Ashburner J. 2009. Computational anatomy with the SPM software. Magn Reson Imaging. doi:10.1016/j.mri.2009.01.006

Barch DM, Burgess GC, Harms MP, Petersen SE, Schlaggar BL, Corbetta M, Glasser MF, Curtiss S, Dixit S, Feldt C, Nolan D, Bryant E, Hartley T, Footer O, Bjork JM, Poldrack R, Smith S, Johansen-Berg H, Snyder AZ, Van Essen DC. 2013. Function in the human connectome: Task-fMRI and individual differences in behavior. Neuroimage 80:169–189. doi:10.1016/j.neuroimage.2013.05.033

Boekel W, Wagenmakers EJ, Belay L, Verhagen J, Brown S, Forstmann BU. 2015. A purely confirmatory replication study of structural brain-behavior correlations. Cortex 66:115–133. doi:10.1016/j.cortex.2014.11.019

Button KS, Ioannidis JPA, Mokrysz C, Nosek BA, Flint J, Robinson ESJ, Munafò MR. 2013. Power failure: why small sample size undermines the reliability of neuroscience. Nat Rev Neurosci 14:365–76. doi:10.1038/nrn3475

Camerer CF, Dreber A, Holzmeister F, Ho T-H, Huber J, Johannesson M, Kirchler M, Nave G, Nosek BA, Pfeiffer T, Altmejd A, Buttrick N, Chan T, Chen Y, Forsell E, Gampa A, Heikensten E, Hummer L, Imai T, Isaksson S, Manfredi D, Rose J, Wagenmakers E-J, Wu H. 2018. Evaluating the replicability of social science experiments in Nature and Science between 2010 and 2015. Nat Hum Behav 1. doi:10.1038/s41562-018-0399-z

Campbell MC, Koller JM, Snyder AZ, Buddhala C, Kotzbauer PT, Perlmutter JS. 2015. CSF proteins and resting-state functional connectivity in Parkinson disease. Neurology 84:2413–2421. doi:10.1212/WNL.0000000000001681

Chambers CD, Dienes Z, McIntosh RD, Rotshtein P, Willmes K. 2015. Registered Reports: Realigning incentives in scientific publishing. Cortex. doi:10.1016/j.cortex.2015.03.022

Choi YY, Shamosh NA, Cho SH, DeYoung CG, Lee MJ, Lee J-M, Kim SI, Cho Z-H, Kim K, Gray JR, Lee KH. 2008. Multiple Bases of Human Intelligence Revealed by Cortical Thickness and Neural Activation. J Neurosci 28:10323–10329. doi:10.1523/JNEUROSCI.3259-08.2008

Cremers HR, Wager TD, Yarkoni T. 2017. The relation between statistical power and inference in fMRI. PLoS One 12:1–20. doi:10.1371/journal.pone.0184923

Desikan RS, Ségonne F, Fischl B, Quinn BT, Dickerson BC, Blacker D, Buckner RL, Dale AM, Maguire RP, Hyman BT, Albert MS, Killiany RJ. 2006. An automated labeling system for subdividing the human cerebral cortex on MRI scans into gyral based regions of interest. Neuroimage 31:968–980. doi:10.1016/j.neuroimage.2006.01.021

Dwan K, Gamble C, Williamson PR, Kirkham JJ. 2013. Systematic Review of the Empirical Evidence of Study Publication Bias and Outcome Reporting Bias - An Updated Review. PLoS One. doi:10.1371/journal.pone.0066844

Eickhoff SB, Yeo BTT, Genon S. 2018. Imaging-based parcellations of the human brain. Nat Rev Neurosci. doi:10.1038/s41583-018-0071-7

Eklund A, Nichols TE, Knutsson H. 2016. Cluster failure: Why fMRI inferences for spatial extent have inflated false-positive rates. Proc Natl Acad Sci U S A 113:7900–5. doi:10.1073/pnas.1602413113

Estle SJ, Green L, Myerson J, Holt DD. 2006. Differential effects of amount on temporal and probability discounting of gains and losses. Mem Cogn 34:914–928. doi:10.3758/BF03193437

Fischl B, Sereno MI, Tootell RBH, Dale AM. 1999. High-resolution intersubject averaging and a coordinate system for the cortical surface. Hum Brain Mapp 8:272–284. doi:10.1002/(SICI)1097-0193(1999)8:4<272::AID-HBM10>3.0.CO;2-4

Fjell AM, Westlye LT, Grydeland H, Amlien I, Espeseth T, Reinvang I, Raz N, Dale AM, Walhovd KB. 2014. Accelerating cortical thinning: unique to dementia or universal in aging? Cereb Cortex 24:919–34. doi:10.1093/cercor/bhs379

Foster, MSLS ED, Deardorff, MLIS A. 2017. Open Science Framework (OSF). J Med Libr Assoc 105:203. doi:10.5195/jmla.2017.88

Foster NEV, Zatorre RJ. 2010. Cortical structure predicts success in performing musical transformation judgments. Neuroimage 53:26–36. doi:10.1016/j.neuroimage.2010.06.042

Franco A, Malhotra N, Simonovits G. 2014. Publication bias in the social sciences: Unlocking the file drawer. Science (80-) 345:1502–1505. doi:10.1126/science.1255484

Gelman A, Loken E. 2014. The garden of forking paths: Why multiple comparisons can be a problem, even when there is no “fishing expedition” or “p-hacking” and the research hypothesis was posited ahead of time. Psychol Bull 140:1272–1280. doi:dx.doi.org/10.1037/a0037714

Genon S, Reid A, Langner R, Amunts K, Eickhoff SB. 2018. How to Characterize the Function of a Brain Region. Trends Cogn Sci. doi:10.1016/j.tics.2018.01.010

Genon S, Wensing T, Reid A, Hoffstaedter F, Caspers S, Grefkes C, Nickl-Jockschat T, Eickhoff SB. 2017. Searching for behavior relating to grey matter volume in a-priori defined right dorsal premotor regions: Lessons learned. Neuroimage 157:144–156. doi:10.1016/j.neuroimage.2017.05.053

Glasser MF, Sotiropoulos SN, Wilson JA, Coalson TS, Fischl B, Andersson JL, Xu J, Jbabdi S, Webster M, Polimeni JR, Van Essen DC, Jenkinson M, WU-Minn HCP Consortium for the W-MH. 2013. The minimal preprocessing pipelines for the Human Connectome Project. Neuroimage 80:105–24. doi:10.1016/j.neuroimage.2013.04.127

Good CD, Johnsrude IS, Ashburner J, Henson RN, Friston KJ, Frackowiak RS. 2001. A voxel-based morphometric study of ageing in 465 normal adult human brains. Neuroimage 14:21–36. doi:10.1006/nimg.2001.0786

Gorgolewski KJ, Varoquaux G, Rivera G, Schwarz Y, Ghosh SS, Maumet C, Sochat V V., Nichols TE, Poldrack RA, Poline J-B, Yarkoni T, Margulies DS. 2015. NeuroVault.org: a web-based repository for collecting and sharing unthresholded statistical maps of the human brain. Front Neuroinform 9:8. doi:10.3389/fninf.2015.00008

Grabitz CR, Button KS, Munafò MR, Newbury DF, Pernet CR, Thompson PA, Bishop DVM. 2018. Logical and methodological issues affecting genetic studies of humans reported in top neuroscience journals. J Cogn Neurosci 30:25–41. doi:10.1162/jocn_a_01192

Gur RC, Richard J, Hughett P, Calkins ME, Macy L, Bilker WB, Brensinger C, Gur RE. 2010. A cognitive neuroscience-based computerized battery for efficient measurement of individual differences: Standardization and initial construct validation. J Neurosci Methods 187:254–262. doi:10.1016/j.jneumeth.2009.11.017

Hyatt CS, Owens MM, Gray JC, Carter NT, MacKillop J, Sweet LH, Miller JD. 2019. Personality traits share overlapping neuroanatomical correlates with internalizing and externalizing psychopathology. J Abnorm Psychol 128:1–11. doi:10.1037/abn0000391

Jeffreys H. 1961. Theory of probability. Oxford, Uk.: Oxford University Press.

Kanai R, Bahrami B, Roylance R, Rees G. 2012. Online social network size is reflected in human brain structure. Proc R Soc B Biol Sci 279:1327–1334. doi:10.1098/rspb.2011.1959

Kanai R, Feilden T, Firth C, Rees G. 2011. Political orientations are correlated with brain structure in young adults. Curr Biol 21:677–680. doi:10.1016/j.cub.2011.03.017

Karama S, Ad-Dab’bagh Y, Haier RJ, Deary IJ, Lyttelton OC, Lepage C, Evans AC. 2009. Positive association between cognitive ability and cortical thickness in a representative US sample of healthy 6 to 18 year-olds. Intelligence 37:145–155. doi:10.1016/J.INTELL.2008.09.006

Karama S, Colom R, Johnson W, Deary IJ, Haier R, Waber DP, Lepage C, Ganjavi H, Jung R, Evans AC. 2011. Cortical thickness correlates of specific cognitive performance accounted for by the general factor of intelligence in healthy children aged 6 to 18. Neuroimage 55:1443–1453. doi:10.1016/j.neuroimage.2011.01.016

Kharabian Masouleh S, Eickhoff SB, Hoffstaedter F, Genon S. 2019. Empirical examination of the replicability of associations between brain structure and psychological variables. Elife 8. doi:10.7554/elife.43464

Kühn S, Schubert F, Gallinat J. 2011. Structural correlates of trait anxiety: Reduced thickness in medial orbitofrontal cortex accompanied by volume increase in nucleus accumbens. J Affect Disord 134:315–319. doi:10.1016/j.jad.2011.06.003

Lindsay DS. 2015. Replication in Psychological Science. Psychol Sci 26:1827–1832. doi:10.1177/0956797615616374

Loken E, Gelman A. 2017. Measurement error and the replication crisis. Science (80-). doi:10.1126/science.aal3618

Marcus DS, Harwell J, Olsen T, Hodge M, Glasser MF, Prior F, Jenkinson M, Laumann T, Curtiss SW, Van Essen DC. 2011. Informatics and Data Mining Tools and Strategies for the Human Connectome Project. Front Neuroinform 5:4. doi:10.3389/fninf.2011.00004

Martínez K, Madsen SK, Joshi AA, Joshi SH, Román FJ, Villalon-Reina J, Burgaleta M, Karama S, Janssen J, Marinetto E, Desco M, Thompson PM, Colom R. 2015. Reproducibility of brain-cognition relationships using three cortical surface-based protocols: An exhaustive analysis based on cortical thickness. Hum Brain Mapp 36:3227–3245. doi:10.1002/hbm.22843

Menary K, Collins PF, Porter JN, Muetzel R, Olson EA, Kumar V, Steinbach M, Lim KO, Luciana M. 2013. Associations between cortical thickness and general intelligence in children, adolescents and young adults. Intelligence 41:597–606. doi:10.1016/j.intell.2013.07.010

Miller KL, Alfaro-Almagro F, Bangerter NK, Thomas DL, Yacoub E, Xu J, Bartsch AJ, Jbabdi S, Sotiropoulos SN, Andersson JLR, Griffanti L, Douaud G, Okell TW, Weale P, Dragonu I, Garratt S, Hudson S, Collins R, Jenkinson M, Matthews PM, Smith SM. 2016. Multimodal population brain imaging in the UK Biobank prospective epidemiological study. Nat Neurosci 19:1523–1536. doi:10.1038/nn.4393

Munafò MR, Nosek BA, Bishop DVM, Button KS, Chambers CD, Percie Du Sert N, Simonsohn U, Wagenmakers EJ, Ware JJ, Ioannidis JPA. 2017. A manifesto for reproducible science. Nat Hum Behav 1:1–9. doi:10.1038/s41562-016-0021

Myerson J, Green L, Warusawitharana M. 2001. AREA UNDER THE CURVE AS A MEASURE OF DISCOUNTING. J Exp Anal Behav 76:235–243. doi:10.1901/jeab.2001.76-235

Natu VS, Gomez J, Barnett M, Jeska B, Kirilina E, Jaeger C, Zhen Z, Cox S, Weiner KS, Weiskopf N, Grill-Spector K. 2019. Apparent thinning of visual cortex during childhood is associated with myelination, not pruning. PANS. doi:10.1101/368274

Nissen SB, Magidson T, Gross K, Bergstrom CT. 2016. Publication bias and the canonization of false facts. Elife 5:1–19. doi:10.7554/eLife.21451

Noble S, Scheinost D, Constable RT. 2019. Cluster failure or power failure? Evaluating sensitivity in cluster-level inference. Neuroimage 116468. doi:10.1016/J.NEUROIMAGE.2019.116468

Nosek BA, Alter G, Banks GC, Borsboom D, Bowman SD, Breckler SJ, Buck S, Chambers CD, Chin G, Christensen G, Contestabile M, Dafoe A, Eich E, Freese J, Glennerster R, Goroff D, Green DP, Hesse B, Humphreys M, Ishiyama J, Karlan D, Kraut A, Lupia A, Mabry P, Madon TA, Malhotra N, Mayo-Wilson E, McNutt M, Miguel E, Paluck EL, Simonsohn U, Soderberg C, Spellman BA, Turitto J, VandenBos G, Vazire S, Wagenmakers EJ, Wilson R, Yarkoni T. 2015. Promoting an open research culture. Science (80-). doi:10.1126/science.aab2374

Open Science Collaboration OS. 2015. Estimating the reproducibility of psychological science. Science 349:aac4716. doi:10.1126/science.aac4716

Owens MM, Hyatt CS, Gray JC, Carter NT, Mackillop J, Miller JD, Sweet LH. 2019. Cortical morphometry of the five-factor model of personality: Findings from the Human Connectome Project full sample. Soc Cogn Affect Neurosci 14:381–395. doi:10.1093/scan/nsz017

Poldrack RA, Baker CI, Durnez J, Gorgolewski KJ, Matthews PM, Munafò MR, Nichols TE, Poline J-B, Vul E, Yarkoni T. 2017. Scanning the horizon: towards transparent and reproducible neuroimaging research. Nat Rev Neurosci 18:115–126. doi:10.1038/nrn.2016.167

Raz N. 2005. Ageing and the Brain. Encycl Life Sci 1–6. doi:10.1038/npg.els.0004063

Riccelli R, Toschi N, Nigro S, Terracciano A, Passamonti L. 2017a. Surface-based morphometry reveals the neuroanatomical basis of the five-factor model of personality. Soc Cogn Affect Neurosci 12:671–684. doi:10.1093/scan/nsw175

Riccelli R, Toschi N, Nigro S, Terracciano A, Passamonti L. 2017b. Surface-based morphometry reveals the neuroanatomical basis of the five-factor model of personality. Soc Cogn Affect Neurosci nsw175. doi:10.1093/scan/nsw175

Rice K, Redcay E. 2015. Spontaneous mentalizing captures variability in the cortical thickness of social brain regions. Soc Cogn Affect Neurosci 10:327–334. doi:10.1093/scan/nsu081

Savic I, Arver S. 2014. Sex differences in cortical thickness and their possible genetic and sex hormonal underpinnings. Cereb Cortex 24:3246–3257. doi:10.1093/cercor/bht180

Schilling C, Kühn S, Romanowski A, Schubert F, Kathmann N, Gallinat J. 2012. Cortical thickness correlates with impulsiveness in healthy adults. Neuroimage 59:824–830. doi:10.1016/j.neuroimage.2011.07.058

Schmitt JE, Raznahan A, Clasen LS, Wallace GL, Pritikin JN, Lee NR, Giedd JN, Neale MC. 2019. The Dynamic Associations Between Cortical Thickness and General Intelligence are Genetically Mediated. Cereb Cortex. doi:10.1093/cercor/bhz007

Schnack HG, Van Haren NEM, Brouwer RM, Evans A, Durston S, Boomsma DI, Kahn RS, Hulshoff Pol HE. 2015. Changes in thickness and surface area of the human cortex and their relationship with intelligence. Cereb Cortex 25:1608–1617. doi:10.1093/cercor/bht357

Sham PC, Purcell SM. 2014. Statistical power and significance testing in large-scale genetic studies. Nat Rev Genet. doi:10.1038/nrg3706

Smith R, Keramatian K, Christoff K. 2007. Localizing the rostrolateral prefrontal cortex at the individual level. Neuroimage 36:1387–1396. doi:10.1016/j.neuroimage.2007.04.032

Smith SM, Nichols TE. 2009. Threshold-free cluster enhancement: Addressing problems of smoothing, threshold dependence and localisation in cluster inference. Neuroimage 44:83–98. doi:10.1016/j.neuroimage.2008.03.061

Van Essen DC, Smith SM, Barch DM, Behrens TEJ, Yacoub E, Ugurbil K. 2013. The WU-Minn Human Connectome Project: An overview. Neuroimage 80:62–79. doi:10.1016/j.neuroimage.2013.05.041

Vul E, Harris C, Winkielman P, Pashler H. 2009. Puzzlingly High Correlations in fMRI Studies of Emotion, Personality, and Social Cognition. Perspect Psychol Sci 4:274–290. doi:10.1111/j.1745-6924.2009.01132.x

Wagenmakers E-J, Verhagen J, Ly A, Bakker M, Lee MD, Matzke D, Rouder JN, Morey RD. 2015. A power fallacy. Behav Res Methods 47:913–917. doi:10.3758/s13428-014-0517-4

Wagenmakers EJ, Wetzels R, Borsboom D, van der Maas HLJ, Kievit RA. 2012. An Agenda for Purely Confirmatory Research. Perspect Psychol Sci 7:632–638. doi:10.1177/1745691612463078

Walhovd KB, Fjell AM, Giedd J, Dale AM, Brown TT. 2017. Through Thick and Thin: a Need to Reconcile Contradictory Results on Trajectories in Human Cortical Development. Cereb Cortex 27:1472–1481. doi:10.1093/cercor/bhv301

Winkler AM, Greve DN, Bjuland KJ, Nichols TE, Sabuncu MR, Håberg AK, Skranes J, Rimol LM. 2018. Joint Analysis of Cortical Area and Thickness as a Replacement for the Analysis of the Volume of the Cerebral Cortex. Cereb Cortex 28:738–749. doi:10.1093/cercor/bhx308

Winkler AM, Kochunov P, Blangero J, Almasy L, Zilles K, Fox PT, Duggirala R, Glahn DC. 2010. Cortical thickness or grey matter volume? The importance of selecting the phenotype for imaging genetics studies. Neuroimage 53:1135–1146. doi:10.1016/j.neuroimage.2009.12.028

