## Supplementary Tables for "Searching for replicable associations between cortical thickness and psychometric variables in healthy adults: empirical facts"

**Table S1.** Distribution of raw phenotypical and behavioral scores in the whole sample.

| **Healthy sample** | **Participants** (n = 420 ; 210 male) |
| --- | --- |
| Age (years)  n-total = 420 | 28.1 $\pm$3.7 (22,37) |
| Education (years)  n-total = 420 | 14.9 $\pm$1.76 (11,17) |
| Anger Affect  n-total = 420 | 48.5 $\pm$8.56 (28,85.4) |
| Anger Aggression  n-total = 420 | 51.9 $\pm$8.96 (43.4,83.1) |
| Anger Hostility  n-total = 420 | 51.12 $\pm$8.8 (36.6, 74) |
| Card Sorting  n-total = 420 | 114.86 $\pm$10.81 (85,143.94) |
| Cognition Early childhood Component  n-total = 420 | 116.8 $\pm$11.05 (85.6 , 153.5) |
| Cognition Crystal Component  n-total = 420 | 117.54 $\pm$9.85 (90.95,147.56) |
| Cognition Fluid Component  n-total = 418 | 115.4 $\pm$12.1 (84.5 , 145.2) |
| Cognition Total Component  n-total = 418 | 121.9 $\pm$15.01 (84.5,153.4) |
| Delay discounting (AUC $200)  n-total = 418 | 0.26 $\pm$0.2 (0.016,0.98) |
| Delay discounting (AUC $40K)  n-total = 418 | 0.5 $\pm$0.3 (0.016,0.98) |
| Dexterity  n-total = 420 | 112 $\pm$10.7 (85,1,148.7) |
| Emotional support  n-total = 420 | 51.2 $\pm$9.6 (15.9,62.5) |
| Emotion recognition (#correct responses)  n-total = 418 | 35.6 $\pm$2.5 (24,40) |
| Emotion recognition (correct responses time)  n-total = 418 | 1833.8 $\pm$349.36 (1250,5020) |
| Handedness  n-total = 420 | 65.3 $\pm$42.7 (-100,100) |
| Visual episodic memory (# correct responses)  n-total = 418 | 35.7 $\pm$2.8 (26,40) |
| Visual episodic memory (correct responses time)  n-total = 418 | 1560.73 $\pm$304.5 (1012.5,3265.25) |
| Language task (Math)  n-total = 412 | 83.6 $\pm$10.15 (51.7,100) |
| Life satisfaction  n-total = 420 | 54.5 $\pm$9.2 (24.2,74.6) |
| List sorting  n-total = 420 | 111.98 $\pm$11.5 (80.8,144.5) |
| Agreeableness  n-total = 418 | 33.1 $\pm$5.9 (10,48) |
| Conscientiousness  n-total = 418 | 34.4 $\pm$5.9 (11,48) |
| Extraversion  n-total = 418 | 30.5 $\pm$6.03 (10,47) |
| Neuroticism  n-total = 418 | 17 $\pm$7.84 (0,43) |
| Openness  n-total = 418 | 28.6 $\pm$6.2 (10,45) |
| PMAT (# correct responses)  n-total = 418 | 16.8 $\pm$4.9 (6,24) |
| Relational task (ACC %)  n-total = 412 | 65.18 $\pm$17.64 (16.7,100) |
| WM Task [2back; Body] (ACC %)  n-total = 418 | 76.84 $\pm$14.3 (18.75,100) |
| WM Task [2back; Body] (median reaction time)  n-total = 418 | 1082.69 $\pm$169.9 (676.5,1514) |
| WM Task [2back; Face] (ACC %)  n-total = 418 | 89.5 $\pm$10.3 (50,100) |
| WM Task [2back; Face] (median reaction time)  n-total = 418 | 924.9 $\pm$170.86 (582,1545.25) |
| WM Task [2back; Place] (ACC %)  n-total = 418 | 90 $\pm$10.3 (31.25,100) |
| WM Task [2back; Place] (median reaction time)  n-total = 418 | 931.12 $\pm$168.99 (518.25,1490) |
| WM Task [2back; Tool] (ACC %)  n-total = 418 | 84.39 $\pm$11.3 (50,100) |
| WM Task [2back; Tool] (median reaction time)  n-total = 418 | 931.47 $\pm$169.43 (543.5,1656.5) |

Data are mean ± SD (minimum-maximum).

* median (minimum-maximum).

Abbreviations: AUC: Area under the curve; ACC: Accuracy; PMAT: Penn matrix test; WM: Working memory;

**Table S2. Summary of exploratory findings.** For each discovery sample size, the number of clusters in which cortical thickness is positively or negatively associated with the tested psychological score is reported. The number of splits (out of 100) in which the clusters were detected are noted in parentheses (i.e. % of splits with at least one significant cluster [in the respective direction]).

|  | # discovery = 70% # total | | # discovery = 50% # total | | # discovery = 30% # total | |
| --- | --- | --- | --- | --- | --- | --- |
|  | # positively associated clusters (split%) | # negatively associated clusters (split%) | # positively associated clusters (split%) | # negatively associated clusters (split%) | # positively associated clusters (split%) | # negatively associated clusters (split%) |
| Anger Affect  n-total = 420 | 6 (3%) | 0 | 12 (4%) | 0 | 5 (3%) | 0 |
| Anger Aggression  n-total = 420 | 26 (13%) | 0 | 14 (10%) | 0 | 28 (13%) | 0 |
| Anger Hostility  n-total = 420 | 6 (4%) | 0 | 1 (1%) | 0 | 10 (6%) | 0 |
| Card Sorting  n-total = 420 | 0 | 0 | 6 (4%) | 0 | 6 (1%) | 0 |
| Cognition Early childhood Component  n-total = 420 | 0 | 0 | 5 (1%) | 3 (2%) | 2 (1%) | 6 (2%) |
| Cognition Crystal Component  n-total = 420 | 0 | 46 (25%) | 0 | 28 (17%) | 0 | 14 (9%) |
| Cognition Fluid Component  n-total = 418 | 0 | 0 | 3 (2%) | 4 (2%) | 5 (2%) | 4 (2%) |
| Cognition Total Component  n-total = 418 | 0 | 10 (3%) | 0 | 7 (5%) | 0 | 12 (4%) |
| Delay discounting (AUC $200)  n-total = 418 | 31 (10%) | 0 | 19 (7%) | 0 | 8 (3%) | 0 |
| Dexterity  n-total = 420 | 3 (2%) | 0 | 2 (1%) | 2 (1%) | 9 (4%) | 2 (1%) |
| Emotional support  n-total = 420 | 1 (1%) | 0 | 11 (5%) | 0 | 13 (4%) | 2 (2%) |
| Emotion recognition (#correct responses)  n-total = 418 | 0 | 0 | 2 (1%) | 0 | 2 (1%) | 0 |
| Emotion recognition (correct responses time)  n-total = 418 | 0 | 0 | 6 (3%) | 0 | 2 (1%) | 2 (1%) |
| Handedness  n-total = 420 | 26 (12%) | 0 | 12 (4%) | 0 | 12 (6%) | 0 |
| Visual episodic memory (# correct responses)  n-total = 418 | 0 | 0 | 0 | 4 (2%) | 0 | 6 (2%) |
| Visual episodic memory (correct responses time)  n-total = 418 | 0 | 4 (2%) | 0 | 0 | 0 | 11 (3%) |
| Language task (Math)  n-total = 412 | 0 | 0 | 6 (2%) | 0 | 3 (1%) | 1 (1%) |
| Life satisfaction  n-total = 420 | 1 (1%) | 0 | 0 | 0 | 0 | 0 |
| List sorting  n-total = 420 | 0 | 4 (3%) | 0 | 11 (7%) | 3 (1%) | 22 (11%) |
| Extraversion  n-total = 418 | 1 (1%) | 0 | 5 (3%) | 0 | 4 (2%) | 0 |
| Neuroticism  n-total = 418 | 46 (12%) | 0 | 24 (14%) | 0 | 11 (5%) | 0 |
| PMAT (# correct responses)  n-total = 418 | 5 (3%) | 0 | 5 (2%) | 0 | 4 (2%) | 0 |
| WM Task [2back; Body] (ACC %)  n-total = 418 | 18 (10%) | 0 | 9 (4%) | 0 | 28 (9%) | 0 |
| WM Task [2back; Body] (median reaction time)  n-total = 418 | 0 | 5 (4%) | 0 | 2 (2%) | 1 (1%) | 11 (3%) |
| WM Task [2back; Face] (ACC %)  n-total = 418 | 15 (7%) | 0 | 15 (6%) | 0 | 13 (4%) | 0 |
| WM Task [2back; Face] (median reaction time)  n-total = 418 | 2 (2%) | 0 | 3 (2%) | 0 | 12 (5%) | 0 |
| WM Task [2back; Place] (ACC %)  n-total = 418 | 0 | 0 | 0 | 0 | 4 (1%) | 0 |
| WM Task [2back; Place] (median reaction time)  n-total = 418 | 0 | 2 (1%) | 0 | 5 (3%) | 1 (1%) | 5 (3%) |
| WM Task [2back; Tool] (ACC %)  n-total = 418 | 0 | 0 | 3 (1%) | 0 | 17 (6%) | 2 (1%) |
| WM Task [2back; Tool] (median reaction time)  n-total = 418 | 1 (1%) | 0 | 8 (4%) | 0 | 11 (5%) | 0 |

Abbreviations: AUC: Area under the curve; ACC: Accuracy; PMAT: Penn matrix test; WM: Working memory;
